# Analyzing microbial evolution through gene and genome phylogenies

**DOI:** 10.1101/2023.08.15.553440

**Authors:** Sarah Teichman, Michael D. Lee, Amy D. Willis

**Affiliations:** Department of Statistics, University of Washington; NASA Ames Research Center and Blue Marble Space Institute of Science; Department of Biostatistics, University of Washington

## Abstract

Microbiome scientists critically need modern tools to explore and analyze microbial evolution. Often this involves studying the evolution of microbial genomes as a whole. However, different genes in a single genome can be subject to different evolutionary pressures, which can result in distinct gene-level evolutionary histories. To address this challenge, we propose to treat estimated gene-level phylogenies as data objects, and present an interactive method for the analysis of a collection of gene phylogenies. We use a local linear approximation of phylogenetic tree space to visualize estimated gene trees as points in low-dimensional Euclidean space, and address important practical limitations of existing related approaches, allowing an intuitive visualization of complex data objects. We demonstrate the utility of our proposed approach through microbial data analyses, including by identifying outlying gene histories in strains of *Prevotella*, and by contrasting *Streptococcus* phylogenies estimated using different gene sets. Our method is available as an open-source R package, and assists with estimating, visualizing and interacting with a collection of bacterial gene phylogenies. dimension reduction, microbiome, non-Euclidean, statistical genetics, visualization

## 1 Introduction

Scientific interest in microbiomes (communities of microscopic organisms in a given environment) has recently expanded due to the growing understanding of the role of the microbiome in human and environmental health, and in conjunction with the decreasing costs of high-throughput sequencing. Microbiomes are often studied through the lens of microbial evolution, which is essential for appropriately assigning taxonomic labels to organisms, comparing communities, and reconstructing a microbial tree of life (Parks et al., 2018; Hug et al., 2016). However, most evolutionary methods were originally developed to study either multicellular organisms, and are not always applicable to bacterial and archaeal populations (Matsen, 2015). Thus, with the growing interest in studying microbiomes, there is an increasing need for methods designed specifically to study bacterial and archaeal evolution.

Phylogenetic trees are a key estimand in studies of evolution. A phylogenetic tree (or *phylogeny*) organizes entities (e.g., individuals, strains, or species) into a tree that reflects shared ancestry between these entities through both its branching structure (*topology*) and branch lengths. We will consider phylogenetic trees in which these entities are either genes or some representation of genomes. We refer to phylogenies that reflect relationships at the gene level as gene trees and to phylogenies that reflect relationships at the genome-level as phylogenomic trees. When pursuing phylogenomics, most microbial investigations estimate a single phylogeny that summarizes the evolution of a set of organisms (see, e.g., Brown et al. (2015); Parks et al. (2018); Imachi et al. (2020)).

While phylogenomic trees summarize evolution at the level of the organisms’ genomes, individual genes in a microbial genome frequently have different evolutionary histories due to incomplete lineage sorting and/or biological processes including horizontal gene transfer (HGT). This can lead to discrepancies between any individual gene tree and a phylogenomic tree, which has led to controversy over the meaning and utility of a genome-level evolutionary summary for microbes (e.g., Bapteste et al. (2009); Boto (2010); Puigbo et al. (2010)). Controversy aside, it has been noted that “mainstream applications of [genome-level] phylogenetics to […] microbes have typically been with the idea of finding ‘the’ tree of such a collection rather than explicitly exploring divergence between various gene trees” (Matsen, 2015). Exploring the divergence between gene trees is the primary motivation for this work.

In this paper, we take the approach of viewing estimated gene trees as complex data objects that necessitate statistical tools for their analysis, and introduce a method and software for investigating gene-level evolutionary divergence in microbial genomes. We propose first estimating both a phylogenomic tree and individual gene-level phylogenetic trees. We then map this set of trees into Euclidean space and use dimension reduction methods to provide a visual comparison between genes’ evolutionary histories. In contrast to traditional approaches to estimating phylogenomic relationships, which combine prespecified sets of genes to estimate a single tree, our approach uses a more expansive collection of gene sequence data to also estimate individual phylogenies. Our approach complements single-gene analyses and phylogenomic analyses by providing insight into varied mechanisms of evolution in bacterial and archaeal organisms. Specifically, it can be used to identify genes with unusual evolutionary histories. This information can then be used to remove genes deemed to be unusual or problematic from a phylogenomic approach. Additionally, this tool can be used to identify genes with common evolutionary histories and potential anomalies in the gene identification, annotation, alignment, or tree estimation processes.

## 2 Proposal

### 2.1 Overview

We propose an interactive, visualization-based approach to exploring gene-level and phylogenomic trees. While it is extremely challenging to compare large collections of phylogenetic trees directly (e.g., through plots or summary statistics), our approach lets the user visualize a set of gene trees and a phylogenomic tree as points in two dimensions. The interactive elements of our tool allow users to (i) identify genes of interest based on the two-dimensional visualization, (ii) examine the phylogeny of the identified genes, and (iii) potentially exclude any outlying genes from an estimate of the phylogenomic tree. We believe that genes with outlying trees (e.g. those depicted as relatively distant points in the low-dimensional representation) are more likely to have been subject to notably different evolutionary pressures and therefore reflect different data generating processes, or encountered potential problems in their gene identification or alignment processes or tree estimation. Thus, outlying (or *incongruous*) trees reflect genes that the researcher may not want to include when attempting to estimate the genome-level evolutionary history of the organisms under consideration.

We give a brief overview of the approach before describing each component in greater detail. Define *𝒯*_*m*+3_ as the set of edge-weighted phylogenetic trees with *m* + 3 tips, *m≥* 2. For *n* genes and *m* + 3 genomes, we start by estimating *T*_*i*_ *∈𝒯*_*m*+3_, the true (unknown) phylogeny of gene *i*, for genes *i* = 1, …, *n*. We denote the estimate of *T*_*i*_ by 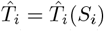 to reflect that this estimator is a function of the aligned gene *i* sequence data, which we denote by *S*_*i*_. We also calculate the tree-valued summary measure 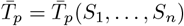 as the phylogenomic tree estimated using gene-level sequence data *S*_1_, …, *S*_*n*_. Once we have our set of *n* gene trees 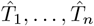 along with 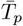, we propose two dimension reduction methods to visualize this set of trees as a scatterplot in ℝ ^*q*^ for small *q* (e.g., *q* = 2 or *q* = 3). Users then can interact with the low dimensional visualization, such as by hovering their cursor over each point on the scatterplot to view the corresponding gene tree, as shown in Figure 1. Users can also investigate the individual gene trees. Given this overview, we now describe each component of the method in greater detail.

**Figure 1.**
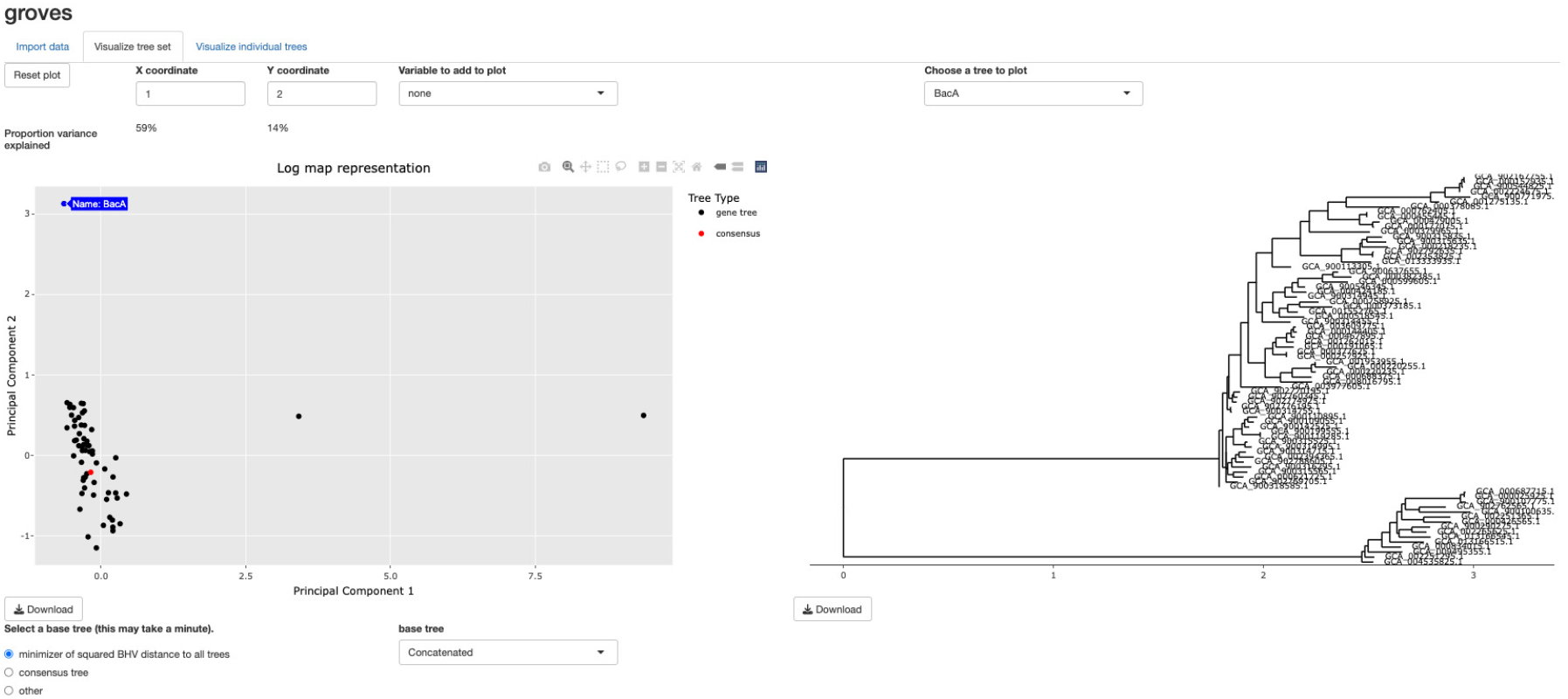
A screenshot of the interactive tool. A scatterplot showing the relationships between a collection of trees (left) can be shown alongside a selected individual gene tree (or collection of individual gene trees) (right). Additional gene-level variables such as functional annotation can also be visualized.

### 2.2 Methodology

Our method begins by estimating the phylogenies of distinct genes in a given collection of microbial genomes and estimating a phylogenomic tree using information from all incorporated genes. These can be estimated via the user’s choice of tree estimator, and the phylogenomic tree does not need to be estimated using the same methodology as the gene-level phylogenies. As is common in studies of microbial evolution (Wu and Eisen, 2008; Segata et al., 2013; Asnicar et al., 2020), in Section 3 we estimate the consensus tree via concatenation. We use the phylogenomics workflow GToTree (Lee, 2019), which by default uses prodigal (Hyatt et al., 2010) to predict genes on input genomes, HMMER3 (Eddy, 2011) to identify target genes, muscle (Edgar, 2021) to align genes, trimal (Capella-Gutiérrez et al., 2009) to trim alignments, and FastTree2 (Price et al., 2010) for approximate maximum-likelihood tree estimation. We use IQTREE2 (Minh et al., 2020) to estimate gene trees via maximum likelihood.

After estimating gene trees, we propose to consider them as objects in the Billera-Holmes-Vogtmann (BHV) space; compute their log maps with respect to a central tree; and then use principal components analysis to visualize them in a low-dimensional Euclidean space. BHV space (Billera et al., 2001), also referred to as “tree space”, is a metric space for the internal branches of phylogenetic trees with the same *m* tips. An *internal branch* is a branch that does not lead to a tip, while a *external branch* does lead to a tip. The BHV distance *γ*(*T*_*i*_, *T*_*i′*_) between two trees accounts for differences in both their topologies (branching structure) and internal branch lengths. Tree space is constructed by representing each possible tree topology by a single non-negative Euclidean orthant, where each coordinate represents the length of an internal branch. The orthants are “glued together” along nearest neighbor interchange topologies. The distance between two trees is defined to be the *L*_2_-length of the shortest possible path between them (see Figure 2). The shortest path between two trees is called the *geodesic*, and is unique (Billera et al., 2001; Owen and Provan, 2011). For two trees with the same topology, the BHV distance between them is simply the *L*_2_-distance between their internal branch lengths. In contrast, for two trees in different topologies, the geodesic path will traverse multiple orthants, and the BHV distance is the sum of the lengths of the linear path segments in each orthant that the geodesic passes through. In this way, the BHV distance encodes information about both branch length differences and topological differences between trees. However, external branch lengths are not encoded in tree space, which we discuss later in this section.

**Figure 2.**
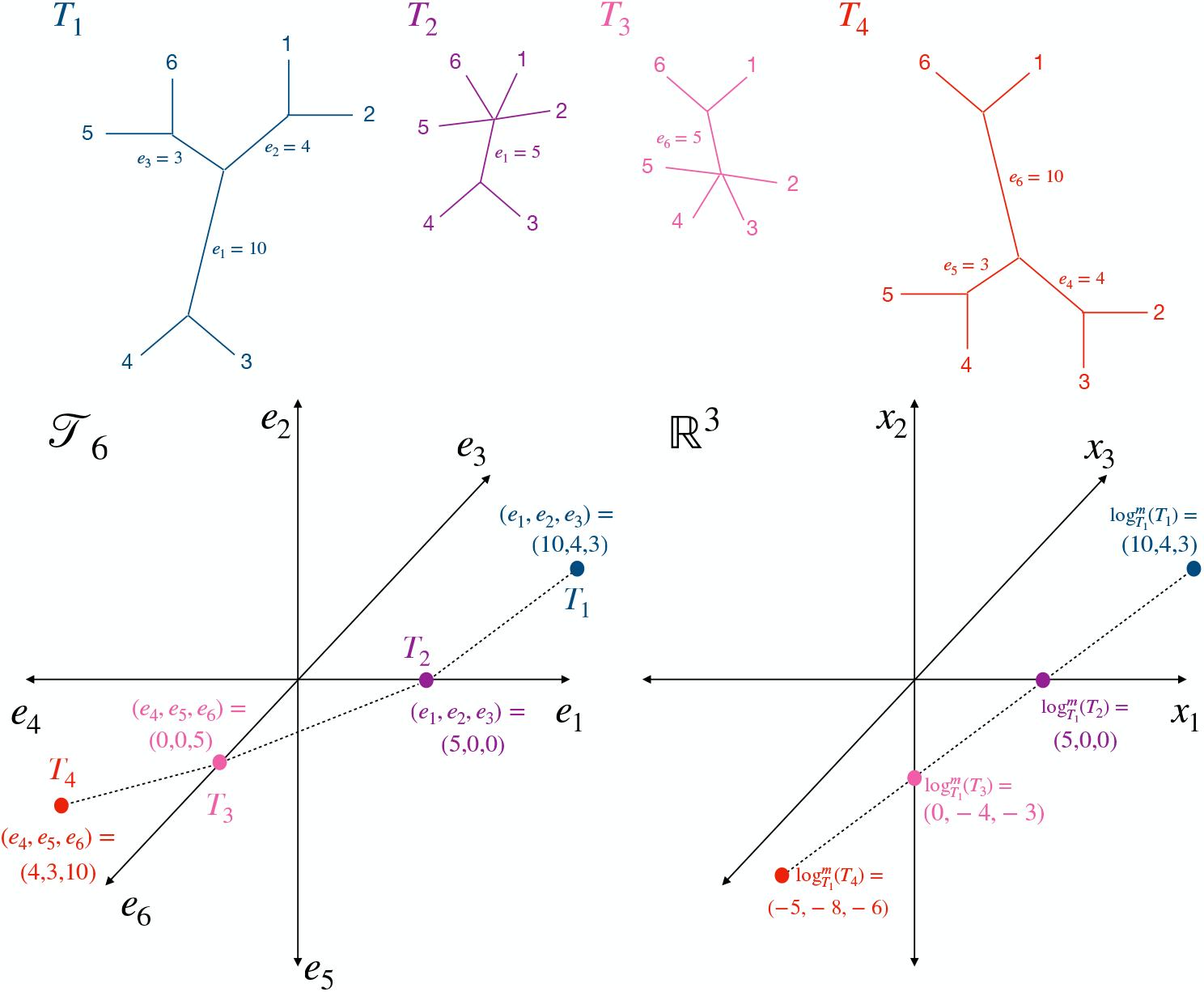
(Top panel) The geodesic path between *T*_1_ and *T*_4_ passes through *T*_2_ and *T*_3_. (Bottom left panel) A representation of the path between *T*_1_ and *T*_4_ in 𝒯_6_, and (bottom right panel) a mapping of 𝒯_6_ around *T*_1_ in ℝ^3^ via the modified log map. Each binary tree topology corresponds to a single non-negative orthant in ℝ^3^, and the orthants are joined along axes corresponding to common branches.

Because of its complex combinatorial structure, it is challenging to visualize trees in their native BHV space. Instead, we find a local Euclidean approximation to BHV space around a central tree, then rely on Euclidean analysis tools such as principal coordinates analysis and scatterplots. To do this, we consider the log map (Barden et al., 2018), which is a mapping from 𝒯_*m*+3_ to ℝ ^*m*^ that captures both geodesic distance and local direction around a base tree. Barden et al. (2018) *define the log map of a tree T* from a base tree *T* ^*\**^ as log_*T \**_ (*T*) = *γ*(*T* ^*\**^, *T*)**v**_*T \**_ (*T*), in which *γ*(*T* ^*\**^, *T*) is the geodesic distance between the two trees, and **v**_*T \**_ (*T*) is a unit vector that represents the direction of the first linear segment of the geodesic path from *T* ^*\**^ to *T*. The modified log map (see Figure 2) can be defined similarly, but as 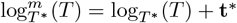, where **t**^*\**^ is the coordinate in ℝ^*m*^ that encodes the internal branch lengths of the base tree (Willis, 2019).

While the log map and modified log map contain information about the topology of a tree and its internal branches, they do not contain information about the external branches. However, external branch lengths provide information about evolutionary distance between entities, and we therefore believe that external branch information should be reflected in our visualizations. We therefore also define the augmented log map, 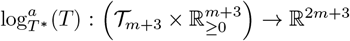, in which the first *m* coordinates are given by the modified log map and the next *m* + 3 coordinates are the lengths of the external branches in a predefined order (e.g., based on alphabetical ordering of the leaf labels). Thus, the augmented log map allows us to embed a collection of phylogenetic trees on *m* + 3 tips into Euclidean space in a way that preserves geodesic distances and local directions from a chosen base tree, as well as external branch lengths.

Having defined the necessary mathematical infrastructure to describe our mapping from tree space to Euclidean space, we are now able to describe our method. We propose to take the augmented log map of 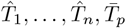 to obtain *n* + 1 vectors in ℝ ^2*m*+3^ representing each estimated gene tree and the phylogenomic tree. We propose two options for choosing the base tree of the augmented log map. The first is to choose the base tree as the tree *t* ∈ **T** that minimizes Σ_*s*∈ **T**_ *γ*(*s, t*)^2^ where **T** is the subset of trees 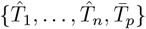 that are binary; that is, have no internal branches of length zero. This will select the most central binary tree in our dataset as the base tree.

An alternative approach to selecting the base tree *t* is to set 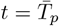; that is, to use the phylogenomic tree as the base tree when computing log maps. While this tree might not be the most central (with respect to squared BHV distance), choosing 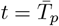 can highlight the differences between the phylogenomic tree and estimated gene trees. This can be especially useful when investigating whether the phylogenomic tree differs substantially from most gene trees, which may occur in situations when the phylogenomic tree estimation process is variable. This often occurs when the tree estimation procedure inexhaustively or stochastically explores tree space, and may be particularly an issue when using maximum likelihood for tree estimation when *m* is large. We provide an example and discussion of this behavior in Appendix Section 3, along with a discussion of the advantages and disadvantages of our two proposed base tree options. Note, however, that our implementation makes it easy to toggle between different selections of the base tree. We suggest that users investigate the sensitivity of the visualization to the base tree (as we illustrate in Appendix Sections 1.1 and 2.1).

After finding 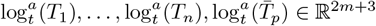 for a given choice of *t*, we perform dimension reduction to create a low-dimensional representation of the set of trees that reflects both the trees’ topologies and branch lengths. Our tool implements principal components analysis (PCA) of augmented log map vectors for dimension reduction. In our data examples shown in Section 3, we use the first two principal coordinates of our *n* + 1 2*m* + 3-dimensional vectors to visualize our tree-valued data objects.

An advantage of our BHV space-based approach is that it enables the exploration of variation in phylogenies with respect to differences in tree topologies and branch lengths. Even for users who are primarily interested in comparing trees topologically, we believe that branch lengths provide information about both the extent of evolution as well as the uncertainty in tree estimation. For example, long branches represent alignments with more bases or amino acids that support a given split in the tree, which corresponds to relatively low uncertainty in the presence of that split in the tree. Correspondingly, shorter branches represent splits with fewer divergent sites. Thus, even for researchers who are primarily interested in exploring differences in tree topologies, our proposed approach still provides important information about uncertainty in topology via the magnitudes of branch lengths. This interplay between topology and branch lengths is a key advantage of BHV space-based analysis: a branch length shrinking to zero is equivalent to moving towards a topological change.

While the above describes our proposed method, we also permit a number of modifications that provide more flexibility for users. For example, we allow the option to construct a multidimensional scaling (MDS) visualization using either BHV distances or Robinson-Foulds (RF) distances between trees. MDS with the RF distance provides a visualization method for researchers who are interested in comparing trees based on topology alone (without regard for branch lengths). In addition, we provide various options for MDS, including both metric MDS and nonmetric MDS. Our software implementation is easily extensible, allowing for further expansion of the possible variations of our general methodology. In practice we find that our proposed approach is highly interpretable, producing scatterplot visualizations where most trees are distributed in all directions around the base tree. This distribution of points in the scatterplot is intuitive and allows for straightforward visual diagnosis of outliers.

### 2.3 Relationship to other work

While a number of studies and tools also make use of low-dimensional representations of a set of trees for analysis, we believe that the novelty of our proposal is three-fold. Firstly, while other methods compute distances between trees and use MDS for dimension reduction (Amenta and Klingner, 2002; Holmes, 2006; Chakerian and Holmes, 2012; Kendall and Colijn, 2016; Gori et al., 2016; Huang et al., 2016; Jombart et al., 2017), we use the log map to find an approximation in Euclidean space for each tree and use PCA for dimension reduction. We believe that PCA has many advantages not shared by MDS, which scales poorly when the number of trees is large because all pairwise distances between trees must be computed. In addition, PCA has advantages for reproducibility when adding new trees to a visualization or comparing across different studies (Willis and Bell, 2018).

Secondly, our approach addresses a key practical limitation of the related work of Willis and Bell (2018), who also proposed PCA on log map-transformed gene trees. Willis and Bell (2018) proposed to use the Fréchet mean of the gene trees as the base tree for computing log maps. Unfortunately, Fréchet means of microbial gene trees are almost always non-binary, including in both examples considered in Section 3. Non-binary trees fall in low-dimensional strata of BHV tree space, resulting in additional distortion and compression of tree space in comparison to log map projections from binary base trees. Therefore, to address the limitation of non-binary Fréchet mean trees, we proposed two alternative options as base trees: the phylogenomic tree, or the binary tree that minimizes the sum of squared BHV distances. The minimization approach is necessarily binary by construction, and we have never seen an example of a non-binary phylogenomic tree. The latter can be explained by the prevalence of concatenation for microbial phylogenomics, which results in many genetic sites for tree estimation and thus the identification of distinct lineage events. Thus, our proposal addresses a prohibitive limitation of the Willis and Bell (2018) methodology, making it applicable to a wider variety of datasets while retaining the advantages of a BHV-based analysis.

Finally, while a number of tree-based visualization methods have analyzed eukaryotic evolution (Hillis et al., 2005; Nye, 2011; Kendall and Colijn, 2016; Gori et al., 2016; Willis and Bell, 2018), our focus on bacterial and archaeal gene trees supports the continuing expansion of microbiome research. Bacterial and archaeal gene trees typically display a greater degree of incongruity than eukaryotic gene trees due to the comparatively high frequency of horizontal gene transfer in these domains. Our emphasis on detecting gene-level phylogeny outliers reflects the greater variability in the distribution of gene trees of bacteria and archaea. Unlike other modern tools for viewing a collection of trees (Huang et al., 2016; Jombart et al., 2017), our tool lets the user choose a tree in the low-dimensional visualization and see its estimated phylogeny, providing rapid insight into why certain trees might appear as outliers. By visually identifying the phylogenomic tree in the two-dimensional visualization and supporting gene selection for phylogenomic tree estimation, our tool is well-suited to improving modern microbial phylogenetic estimation as it is currently practiced.

## 3 Data Analyses

We envisage three primary use cases for our exploratory method. We briefly describe these settings before illustrating the approach on bacterial genome datasets.

- Identifying gene-level phylogeny outliers: Any individual gene tree can differ from the remaining trees in the collection through its topology, branch lengths, or both. Our method can be used to identify outlying gene trees via the low-dimensional visualization, then explore more deeply by plotting the trees itself. This approach can be used to generate hypotheses about drift-based or selective evolution on specific microbial traits, or help identify genes that may have undergone HGT. In Section 3.1 we analyze *Prevotella* genomes and identify and interrogate three genes that are outlying in their phylogenies.
- Contrasting multiple sets of genes for phylogenomics: In microbiome data analysis, phylogenetic trees are commonly estimated by concatenating alignments of multiple genes. However, it is not common practice to investigate the robustness of the estimated tree to the choice of genes that are input to the alignment. Our tool streamlines investigating robustness by easily allowing the removal of genes from a concatenated alignment then re-estimating the phylogenomic tree. We illustrate this approach by comparing the phylogenomic tree estimated using only ribosomal genes for *Streptococcus* genomes to a tree estimated using a fuller set of single-copy orthologous genes in Section 3.2.
- Highlighting anomalies in preprocessing: Finally, our tool can flag potential issues with gene identification, sequence alignments, and tree estimation. For example, an outlying tree may be caused by a tree estimation algorithm that failed to converge, or an issue performing the multiple sequence alignment. We provide a brief example of this use case in Section 4.

For full details on software and database versions used in this section, please see Supplementary File `software-version.csv`.

### 3.1 Prevotella

*Prevotella* is a genus of gram-negative anaerobic bacteria that are present in the human gut, oral, and vaginal microbiomes. The abundance of *Prevotella* in the human gut microbiome is associated with geography, lifestyle, and diet (Tett et al., 2021). Understanding the relatedness of different strains within this genus is essential in studying concepts such as patterns of ecological niches, genomic elasticity, diversification, and host-microbe-phage interactions.

We wish to estimate a phylogenomic tree for the *Prevotella* genus, to explore gene-level histories of *Prevotella* genes, and to understand the concordance between the gene-level and phylogenomic trees. In order to have broad representation across the *Prevotella* genus and work with high quality genomes, we consider the 383 species representatives in the Genome Taxonomy Database (GTDB) (Parks et al., 2018, 2020). To develop a gene set, we first use HMMER (Eddy, 2011) to identify all protein families from the Pfam database (Mistry et al., 2021) in our *Prevotella* genomes that are present in a single copy in a subset of the genomes. We order the Pfams (genes) by their prevalence in the *Prevotella* genomes, omitting those in the lowest 10% of prevalence, resulting in 63 genes for analysis. We then consider only genomes for which all of these genes are present, leaving us with 78 genomes and 63 genes. These choices result in a set of genomes that each contain all of our genes of interest, covering much of the breadth of the *Prevotella* genus as well as a set of genes that are broadly prevalent across the genus. Furthermore, manual comparison of 63 trees with 78 tips would be intractable, motivating our streamlined method for their analysis.

After constructing our gene and genome set, we build gene level and concatenated alignments using GToTree (Lee, 2019), which uses prodigal (Hyatt et al., 2010) to predict genes on input genomes, HMMER3 (Eddy, 2011) to identify target genes, muscle (Edgar, 2021) to align genes, and trimal (Capella-Gutiérrez et al., 2009) to trim alignments. We use IQ-Tree (Minh et al., 2020) to estimate 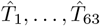 and 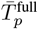, where 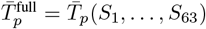. We then construct the visualization described in Section 2.2, choosing the minimal BHV-distance binary tree as the base tree for our log maps. Interestingly, for this analysis, this tree is 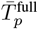, and thus, our two options for choosing the base tree (the phylogenomic tree or the binary tree that minimizes the mean of squared BHV distances) coincide in this instance. Additionally, the unresolved Fréchet mean tree for this set of trees has a BHV distance from the phylogenomic tree of 0.372, which is the smallest distance between any pair of trees in the dataset. A static rendering of our visualization is shown in Figure 3. The first principal component explains 59% of the variance in the log map vectors for the set of trees, and the second principal component explains 14%. Because the log map transformation can map multiple trees in BHV space to the same vector, there is variation between trees in tree space that is not captured in the log map vectors. Therefore, the proportion of variance of the log map vectors explained by each principal component will in general overstate the tree variation explained by each principal component.

**Figure 3.**
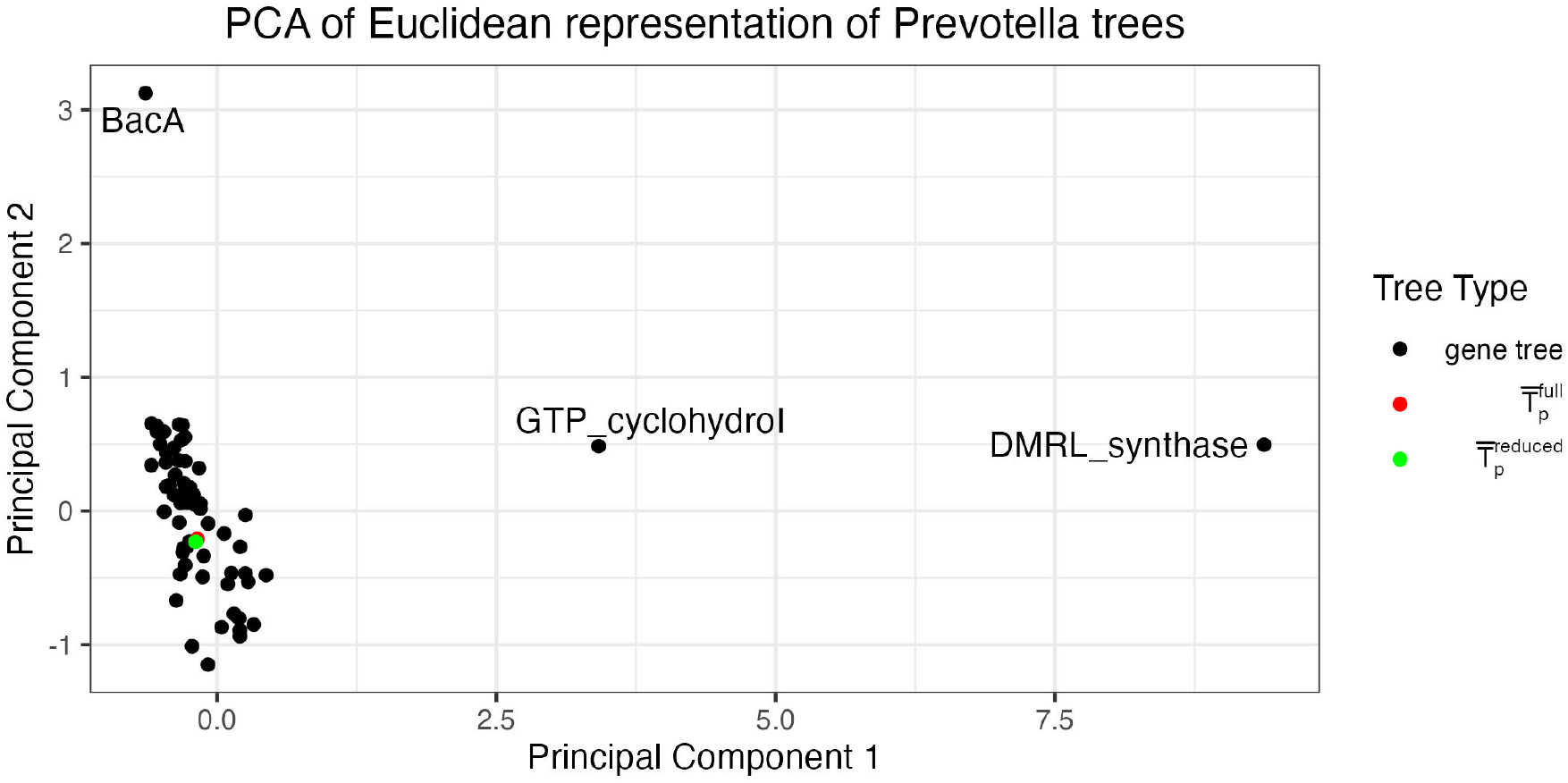
The proposed visualization of 63 gene trees constructed from 78 genomes from the *Prevotella* genus, depicted by a two-dimensional scatterplot. Three visibly outlying gene trees are labeled. A phylogenomic tree constructed from the full gene set is shown in red and a phylogenomic tree constructed from a reduced gene set after removing the three outlying genes is shown in green.

In Figure 3, we clearly observe three genes that have outlying phylogenies relative to the full gene set. Two of these outliers are with respect to the first principal component and correspond to genes DMRL_synthase and GTP_cyclohydroI. The third outlier is with respect to the second principal component and corresponds to the gene BacA. The rest of the gene trees are clustered together in the bottom left of the plot, with 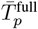 shown in red in the middle of the cluster of gene trees. The placement of 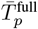 in the middle of the cluster of non-outlying gene trees suggests that there are not major differences in branch lengths nor topology between the concatenated tree and the non-outlying gene trees. To investigate the sensitivity of our estimated phylogenomic tree to the outlying genes, we re-estimated it after removing the three declared outlying genes from the concatenated alignment. We call this tree 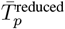, and plot its log map in Figure 3 in green. These two concatenated trees are in almost the same location in the log map visualization, with a RF distance of 14 and a BHV distance of 0.15 (which are the smallest RF and BHV distances between any pairs of trees in this analysis), suggesting that the initial phylogenomic tree is robust (on the scale of topological variation present in the dataset) to the inclusion of the outlying DMRL_synthase, GTP_cyclohydroI and BacA genes. Contrasts with visualization via MDS can be found in Appendix Section 1.2, and visualization via t-SNE and UMAP can be found in Appendix Section 1.3.

We now investigate the outlying gene trees further. The three gene tree outliers along with 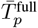are shown in Figures 4a–4d. All trees are visualized using ggtree (Yu et al., 2017). For ease of viewing, we replace tip labels with numeric labels (see Supplementary File `Prevotella-data.csv` for a key to match labels with accession numbers). We see from Figures 4a and 4b that both of these estimated gene trees have a single long external branch leading to tip 51, which corresponds to a genome classified as *Prevotellaceae* bacterium UBA4332 (GCA_900316295.1). This branch length has a first principal component loading of 1.00, relative to a total sum of loading magnitudes of 1.51 across all log map features, giving it a relative weight of 0.66. Our visualization tool provides us an easy way to identify that this long branch is common to these two gene trees and substantially large in comparison with the branches in other gene trees and the phylogenomic trees. This genome (GCA_900316295.1) could be investigated further to understand why DMRL_synthase and GTO_cyclohydroI seem to have distinct evolutionary histories as compared to the rest of the incorporated genes. Visualizations of the alignments for these two genes can be seen in the Supplementary Material (alignment visualizations are generated with NCBI’s Multiple Sequence Alignment Viewer (Sayers et al., 2019)).

**Figure 4.**
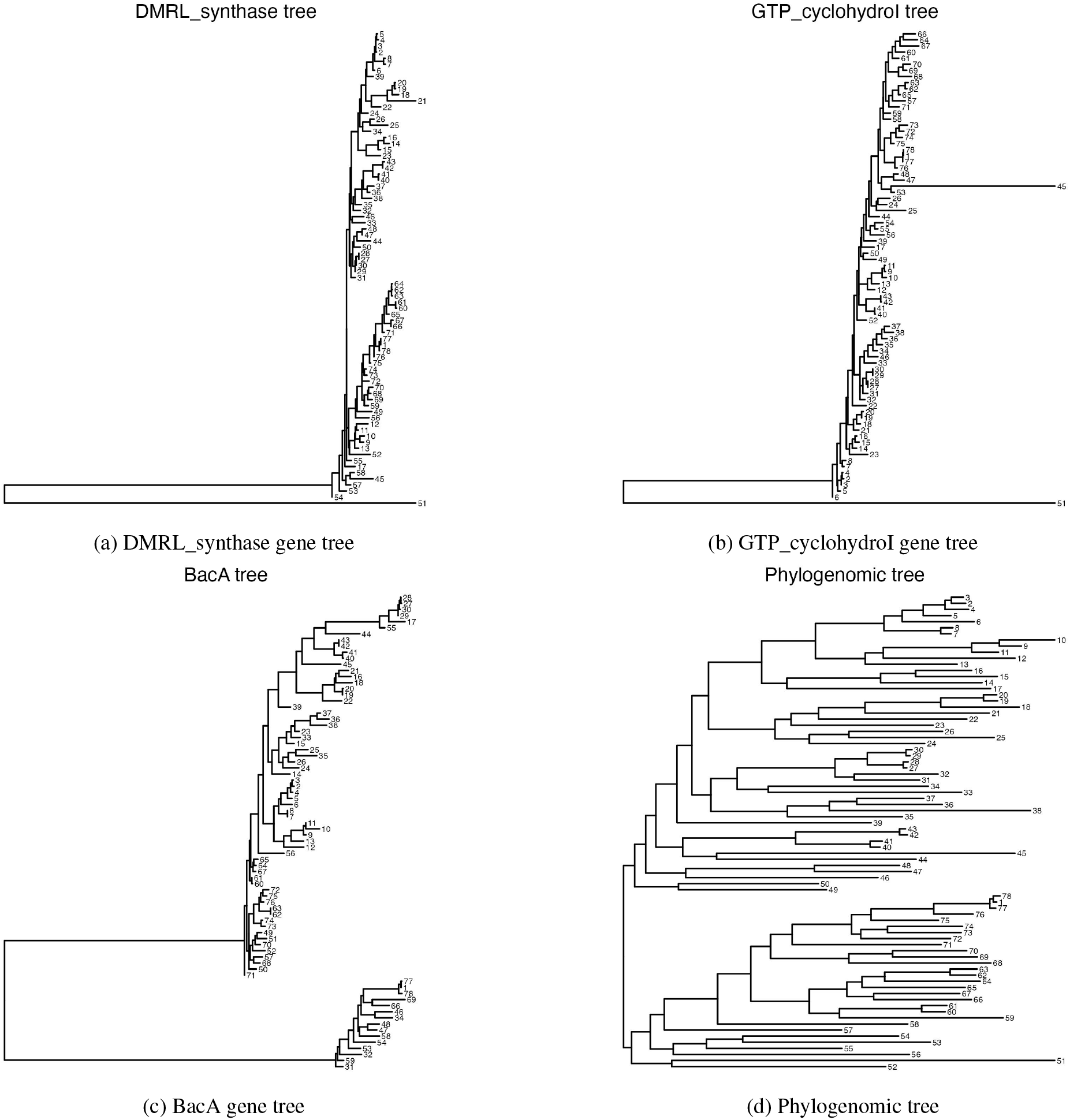
The estimated gene trees for the three outlying *Prevotella* genes identified in Figure 3, as well as the estimated phylogenomic tree (bottom right). All trees are rooted at their mid-point. The DMRL_synthase (top left) and GTP_cyclohydroI (top right) gene trees both include one long branch leading to tip 51. The BacA (bottom left) gene tree includes one long branch separating two clades of tips. No strikingly long branches are present in the phylogenomic tree.

Similarly, the second principal component in our plot differentiates the gene BacA from the rest of the estimated gene and phylogenomic trees. In Figure 4c we observe one long internal branch that separates this gene in 15 genomes from the gene in the remaining 63, while in the phylogenomic tree these 15 are spread across the two major clades (Figure 4d). Furthermore, gene-level differences in each of these genomes can be clearly observed in the amino acid alignments, which can also be seen in the Supplementary Material. The supplementary file `Prevotella-data.csv` holds additional information for the incorporated genomes (e.g., source of the genome), but no clear trends emerge that might contribute to this gene-level phylogeny. It is possible this particular gene may be commonly transferred horizontally. Ultimately, the easy identification of these genes by our approach motivates further exploration into their sequence divergence for the two sets of genomes that are depicted by the large clades in Figure 4c. We additionally investigate the concordance between 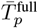 and the gene trees, which can be found in Appendix Section 1.4. We observe the “tree of tips” phenomenon described by Thiergart et al. (2014), in which splits near the tips on the tree have higher gene tree support and splits near the center of the tree have lower support.

### 3.2 Streptococcus

The above analysis provided an example of how to use our proposed method for generating hypotheses about differences in evolutionary histories between genes. We now illustrate how to assess the robustness of the estimated phylogenomic tree to different gene sets. In this analysis, we investigate the differences in phylogenomic trees built from a large set of genes compared to a smaller set of functionally specific (ribosomal) genes. We consider the genus *Streptococcus*, which is of particular relevance in public health and medicine because of its potential pathogenicity.

Similar to Section 3.1, we consider all 301 GTDB representative species genomes for *Streptococcus*; identify 196 protein families (genes) that appear in a single copy in more than 90% of the representative genomes; and analyze the 106 genomes that contain all 196 genes of interest. This large set of genes and genomes provides a good test of our tool because trees with 106 tips are difficult to compare visually, especially a set of 196 trees of this size. Out of the 196 genes that we identified, 38 are ribosomal genes. In general, ribosomal genes are present in bacterial genomes in single copy and are considered essential “core” genes. As a result, they are commonly chosen as an appropriate gene set for constructing a concatenated alignment. We are interested in comparing the phylogenomic tree estimated using the 38 ribosomal genes with the phylogenomic tree estimated with the full set of 196 genes, which represent a wider range of genomic functions. We use GToTree and IQ-Tree to estimate 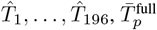, and 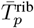, where 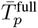 is built from the full set of 196 genes and 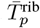 from the set of 38 ribosomal genes.

A static output of our tool is shown in Figure 5a. Interestingly, the phylogenomic tree is again the tree in the set with the minimum mean squared BHV distance from the other trees. This tree has a BHV distance from the unresolved Fréchet mean tree of 0.138, which is less than 97% of the BHV distances between trees in the dataset. The first principal component explains 44% of the variance in the log map vectors for the set of trees and the second principal component explains 10%. We observe a tight cluster of ribosomal gene trees and a more distributed cloud of non-ribosomal trees, suggesting that the estimated ribosomal gene trees are generally more similar to each other than to the other estimated gene trees. Due to relatively high purifying selective pressures, ribosomal proteins tend to evolve more slowly than the majority of genes. Therefore we generally expect their phylogenies to have shorter branch lengths than genes that are less functionally constrained. However, this pattern is not the case for every ribosomal gene, and we observe two ribosomal genes (Ribosomal_S30AE and Ribosomal_L9_C) that are further from the ribosomal gene tree cluster with respect to the first principal component. While these appear to be outlying amongst the ribosomal genes, they are not unusual when compared to the entire collection of gene trees. We also observe two non-ribosomal gene trees, DUF1934 and DUF3270, which appear as potential outliers amongst all genes.

**Figure 5.**
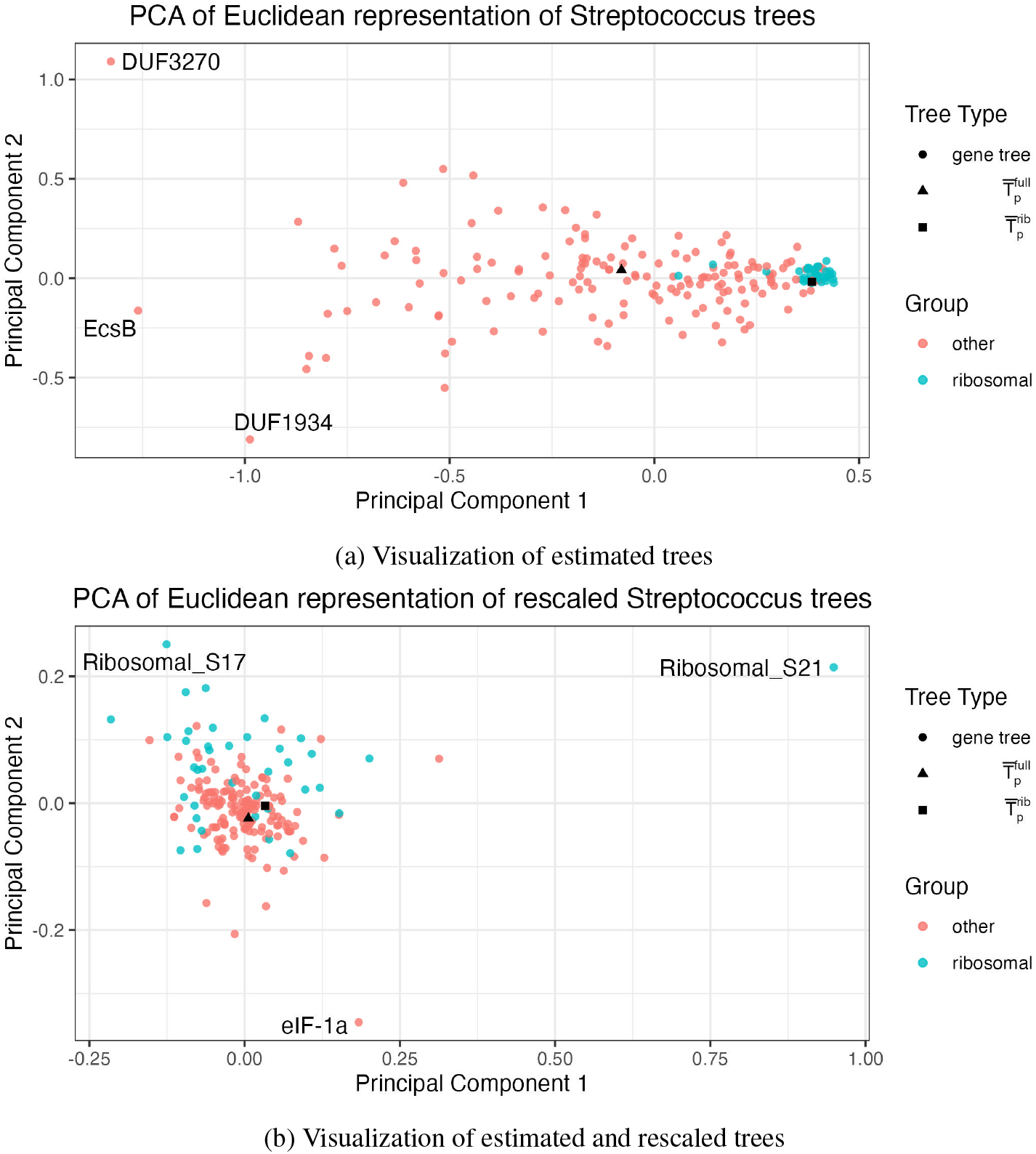
The proposed visualization of 196 gene trees estimated from 106 genomes in the *Streptococcus* genus shown as a two-dimensional scatterplot (upper). The same visualization is shown after rescaling the trees (lower), where the rescaling is performed by dividing all branch lengths on each tree by the sum of the branch lengths for that tree. A phylogenomic tree constructed from the full gene set is shown as a black triangle and a phylogenomic tree constructed from a subset of ribosomal genes is shown as a black square. Gene trees are colored by whether or not they are ribosomal genes.

We can also see a noticeable difference between 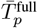 and 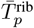 in Figure 5a. In order to compare 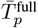 and 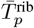 we can compare their distances to distances among other pairs of trees. The BHV distance between the two phylogenomic trees is 0.43, which is smaller than 75% of the BHV distances between gene trees and phylogenomic trees in this analysis. However, the RF distance (based solely on topology) between the two phylogenomic trees is 140, which is smaller than 99.9% of distances between pairs of trees in the analysis. Finally, the difference in the sum of squared branch lengths between the two phylogenomic trees is 2.93, which is smaller than 43% of the distances between trees in the analysis. In general, the estimated ribosomal tree had shorter branches compared to the full gene set tree (ribosomal vs. full tree branch length mean: 0.003 vs. 0.017; median: 0.0008 vs. 0.0090). Taken together, this suggests that these two phylogenomic trees are very similar in their topologies, but differ in their branch lengths. We conclude that the separation implied by the log map visualization is most likely driven by differences in branch lengths, not topology, between these two trees.

Motivated by our observation that branch lengths likely drive the difference between the ribosomal and full phylogenomic trees, we investigate the impact of rescaling *all* trees’ branch lengths on the visualization. We rescale each tree by dividing each individual branch length by the sum of the branch lengths on the given tree. This modification means that our visualization highlights differences between trees with respect to topology and relative branch lengths, but not absolute branch lengths. This is also an option within the interactive visualization. We show the resulting visualization in Figure 5b. We see that the ribosomal gene trees are much more evenly spread among the other gene trees than in Figure 5a, and that the two phylogenomic trees are visibly closer. This provides further evidence that much of the differences between the ribosomal and full phylogenomic trees that we observed in Figure 5a can be attributed to differences in branch lengths. This is unsurprising because as mentioned above, ribosomal genes are typically under greater purifying selective pressures. In the rescaled visualization, there remain a few noticeable outliers that could be investigated further: Ribosomal_S21, Ribosomal_S17, and eIF-1a. These trees all have one or two branches that are especially large relative to other branches in the tree. Notably, we did not detect these differences in the original visualization because none of these long branches were long in comparison to branches on other gene trees prior to rescaling. All trees and rescaled trees are available as Supplementary Data, a MDS visualization can be found in Appendix Section 2.2, and t-SNE and UMAP visualizations can be found in Appendix Section 2.3.

## 4 Discussion

In this paper, we introduced an easy-to-use, interactive method for visualizing a set of trees. Our method was motivated by expanding interest in interrogating microbial evolution, specifically in the context of comparing individual gene evolutionary histories, and the relatively few tools available to facilitate such analyses. Our approach is to represent each phylogeny as a vector in ℝ^2*m*+3^ by taking a local linear approximation to tree space around a central tree, then performing dimension reduction (such as PCA) to view each tree in low-dimensional Euclidean space. The approach of performing PCA on tree approximations is notably different from the approach of Nye (2011), in which a version of PCA is performed in tree space to identify sources of variation in a set of trees. This is also distinct from most other tools for visualizing a set of phylogenies, which compute dissimilarities between each tree and perform MDS on the dissimilarity matrix (Daubin et al., 2002; Amenta and Klingner, 2002; Hillis et al., 2005; Holmes, 2006; Chakerian and Holmes, 2012; Kendall and Colijn, 2016; Jombart et al., 2017; Zhu et al., 2019). Our approach also contrasts with the related method of Willis and Bell (2018) by addressing its main limitation of attempting to locally project tree space around the Fréchet mean tree, which commonly lies at the boundary of two orthants and results in a poor local approximation of the metric space. While our method could be applied to the analysis of any collection of genes, it is especially well-suited to microbial datasets because it integrates with common methods for estimating phylogenomic trees via a concatenated alignment, provides tools for investigating the robustness of the phylogenomic tree estimates to the incorporated genes, and is computationally feasible for analyzing all genes that are common to a set of microbial genomes.

Despite substantial advantages in ease-of-use and intuitive interpretations of the visualization, our approach inherits the limitations of BHV tree space. In particular, BHV geodesics and log maps are only defined for trees that share identical leaf sets. As a result, our tool is restricted to analyzing sets of genes and genomes for which all genes are present in all genomes. In practice, we address this by limiting our analysis to genes present in 90% of genomes, then restricting to genomes that contain all of these genes. However, users could use their own approach if they have specific genes or genomes that they would like to include in their analysis. One implication of this is that more genes can be studied when the incorporated genomes are more closely related evolutionarily. In contrast, datasets with taxa that span phyla or domains will have fewer genes common to many genomes, and may result in a small number of genes for analysis. This limitation is shared by many cross-genome comparative (often called *pangenomic*) analyses. While the absence of a gene in a genome may reflect true biological variation, genes may also be absent due to limitations of genome sequencing and reconstruction. Thus, there are many reasons why a microbial gene may be excluded from a given analysis, and the limitations of BHV space in comparing genomes with different gene sets is one possible reason. Ongoing work to reconcile BHV space across trees with different leaf sets (Grindstaff and Owen, 2020; Ren et al., 2017) may ultimately enable local linear approximations of BHV spaces of different leaf sets, and thus more comprehensive comparisons of gene sets, in the future.

The goal of this work is to demonstrate that investigating gene trees as well as summary phylogenomic trees can uncover interesting evolutionary signal and to offer a method to investigate and analyse sets of trees. However, if a user’s goal is instead to classify gene trees or perform another supervised task, other multivariate analysis methods tailored to that task could be used with the log map vectors as input. We provide an example of using the log map vectors for classification in Appendix Section 2.4.

We provide an easy-to-use software implementation of our proposed methodology. This tool enables a visual exploration of gene-level evolutionary differences across sets of trees, which facilitates the identification of genes with unique (or anomalous) evolutionary histories. Our software also allows for the straightforward refinement of genes being used in estimating a phylogenomic tree, including sensitivity analyses for adding or removing genes. As well as implementing our proposed visualization, our software can also be used to perform dimension reduction with MDS, tSNE, or UMAP, and then plot the results. Finally, we provide the option to output the log map vectors for trees used in an analysis, which could then be used as input to software that performs other types of analyses not included in our software package. Our tool is available as both a workflow or shiny app in our open-source R package at github.com/statdivlab/groves. Code to reproduce the data analyses in Section 3 is available at github.com/statdivlab/groves_supplementary.

## Acknowledgments

The authors gratefully acknowledge support from the National Institute of General Medical Sciences (R35 GM133420) and thank the editor and two reviewers for constructive suggestions.

## Code and data availability

Code and data for reproducing the results of this manuscript are available online at github.com/statdivlab/ groves-supplementary. The unprocessed data are available online, as described in Section 3.

