## Appendix for "Analyzing microbial evolution through gene and genome phylogenies"

### Additional data analyses: *Prevotella* and *Streptococcus* gene trees

August 15, 2023

#### 1 Prevotella

##### 1.1 Sensitivity to the choice of log map base tree

The purpose of this additional analysis of the *Prevotella* gene trees is to investigate the sensitivity of the proposed visualization to changes in the base tree used in the log map transformation. Figure 1 shows the proposed visualization constructed using the phylogenomic tree as the base tree (which, in this case, is also the tree with the lowest mean squared BHV distance from the other trees in the set) as well as the visualization constructed with five different base trees. These five trees have the next smallest mean squared BHV distance from other trees in the set after the phylogenomic tree. Each of the visualizations in Figure 1 show the same three outlying genes (unlabeled for visual simplicity): BacA, DMRL_synthase, and GTP_cyclohydroI. There are subtle differences between the visualizations including the spread of the non-outlying trees in the second principal component and the orientation of the BacA gene (the outlier in the second principal component) compared to the rest of the trees. However, overall, the same conclusions can be drawn by each of the visualizations in Figure 1.

In Figure 2, we now show the proposed log map visualization with base trees chosen to be the three outliers visible in Figure 1. The visualizations that use BacA and GTP_cyclohydrI as base trees are very similar to the one that uses the phylogenomic tree as the base tree. Interestingly, when DMRL_synthase is the base tree, BacA is no longer identifiable as a clear outlier in the second principal component.

Our recommendation for choosing base trees is to choose a tree that is central to the tree set (in terms of BHV distance). Such a choice will map as much information as possible from BHV tree space to the log map representation. However, we strongly suggest running a sensitivity analysis (similar to those described above) to investigate if the results of the visualization are relatively robust to the base tree choice.

Finally, we investigate the result of choosing a base tree with the topology drawn uniformly-at-random, i.e., a base tree with a topology uninformed by the gene tree data. While gene trees rarely all have the same topology as each other, they tend to have similar topologies relative to the number of possible tree topologies. Thus, they exist in a small number of BHV orthants. We hypothesize that the proposed visualization was relatively robust to choice of base tree in Figures 1 and 2 because all of the trees in the analysis exist in this small number of nearby orthants in BHV space. By this rationale, choosing a base tree from an orthant that is farther away from the set of trees could distort the visualization, because the direction of the first segment of the geodesic from the base tree to each tree in the analysis will be similar. Figure 3 shows the proposed visualization using a randomly generated tree on 78 tips as the base tree. The topology is drawn at random using the function `rtree` from the package ape [Paradis and Schliep, 2019], and the branch lengths are drawn at random from the empirical distribution of branch lengths from all trees in this *Prevotella* dataset. Results

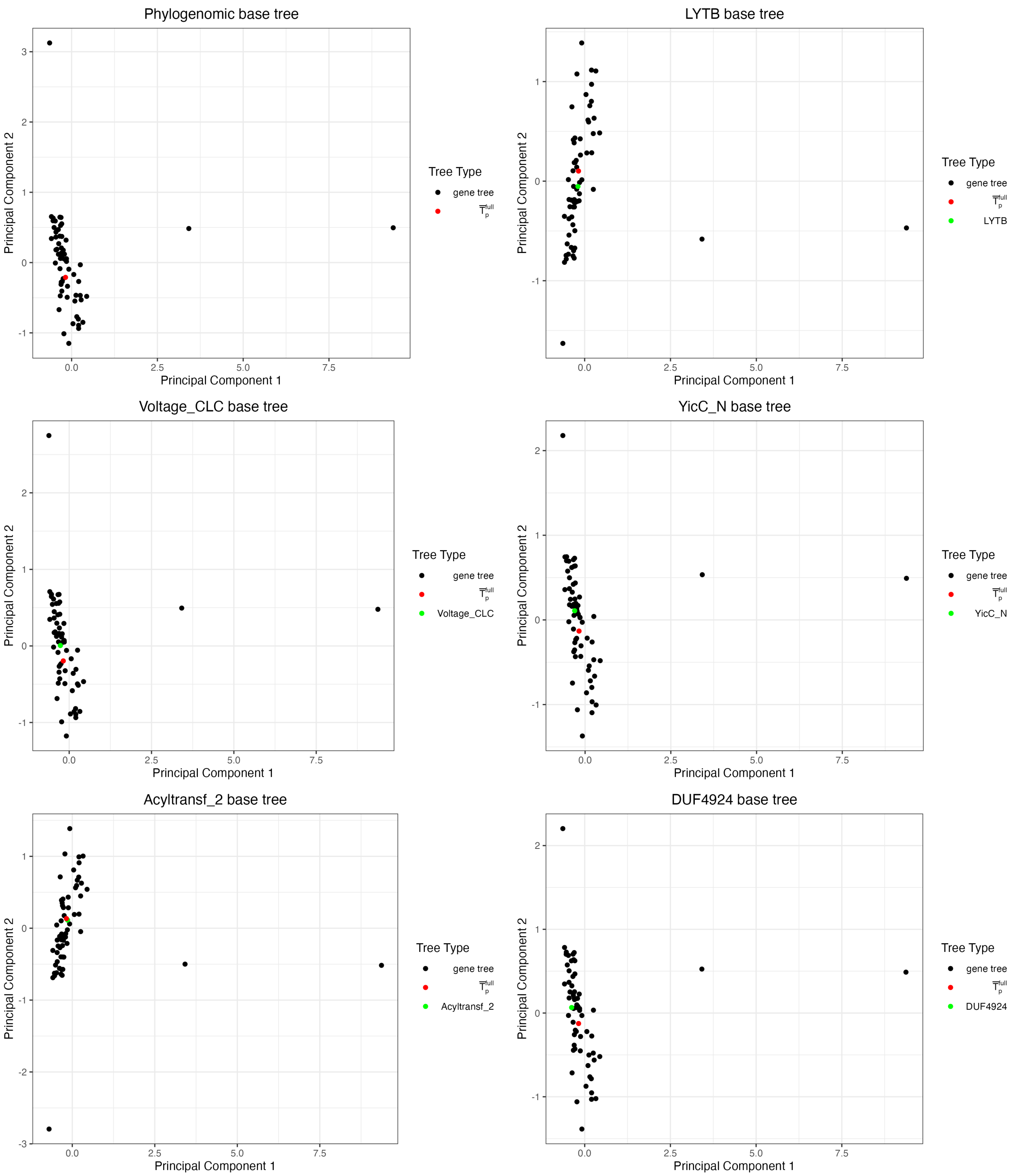


Figure 1: The proposed visualization of 63 gene trees constructed from 78 genomes from the *Prevotella* genus, depicted by a two-dimensional scatterplot. Each subplot is constructed using a different tree as the base tree in the log map (see subplot titles). The five chosen gene trees (LYTB, Voltage_CLC, YicC_N, Acyltransf_2, and DUF4924) have the smallest mean squared BHV distances from other trees in the dataset.

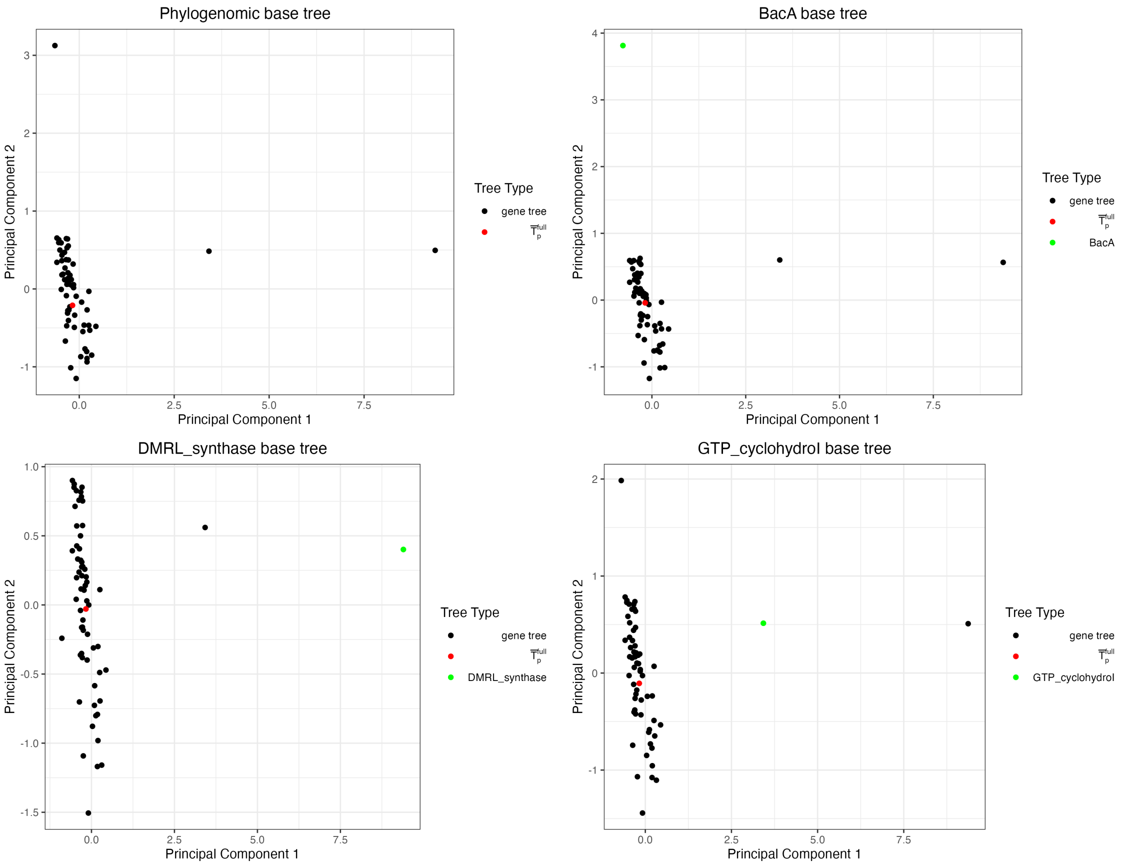


Figure 2: The proposed visualization of 63 gene trees constructed from 78 genomes from the *Prevotella* genus, depicted by a two-dimensional scatterplot. Each subplot is constructed using a different tree as the base tree in the log map. BacA, DMRL_synthase, and GTP_cyclohydroI are the outlying genes identified in the plot created with the phylogenomic tree as the base tree.

look very similar when different random base trees are generated. This plot shown in Figure 3 has the same patterns as the majority of the plots in Figures 1 and 2, with the exception of the randomly generated base tree, which is an outlier in the second principal component. This plot shows that while the random base tree is clearly an outlier from the set of trees in the analysis, the patterns in the plot are even robust to this choice of topology-uninformed base tree.

##### 1.2 MDS visualization

Figure 4 provides a visualization of the *Prevotella* gene trees and phylogenomic tree using metric MDS of BHV distances. This plot shows the same signals as the proposed visualization, with the phylogenomic tree towards the center of a cluster of gene trees, with DMRL_synthase and GTP_cyclohydroI as outliers in the first dimension and BacA as an outlier in the second dimension. However, the cloud of non-outlying trees is less dense in the MDS plot in Figure 4 compared to the plots in Figures 1 and 2. The cloud of points is spread out along both MDS dimensions 1 and 2 in Figure 4, versus only along principal component 2 in the PCA plots.

##### 1.3 tSNE and UMAP visualizations

In addition to the more traditional dimension reduction methods of PCA and MDS, we can also consider performing dimension reduction using t-distributed stochastic neighbor embedding (t-SNE) and uniform manifold approximation and projection (UMAP) on the log map vectors. We implement t-SNE using the function `Rtsne` from the R package of the same name [Krijthe, 2015], and UMAP using the function `umap` from the R package of the same name [Konopka, 2023]. As both of these dimension reduction algorithms are stochastic, we consider visualizations created with four different initial seeds. The t-SNE visualizations can be seen in Figure 5 and the UMAP visualizations in Figure 6.

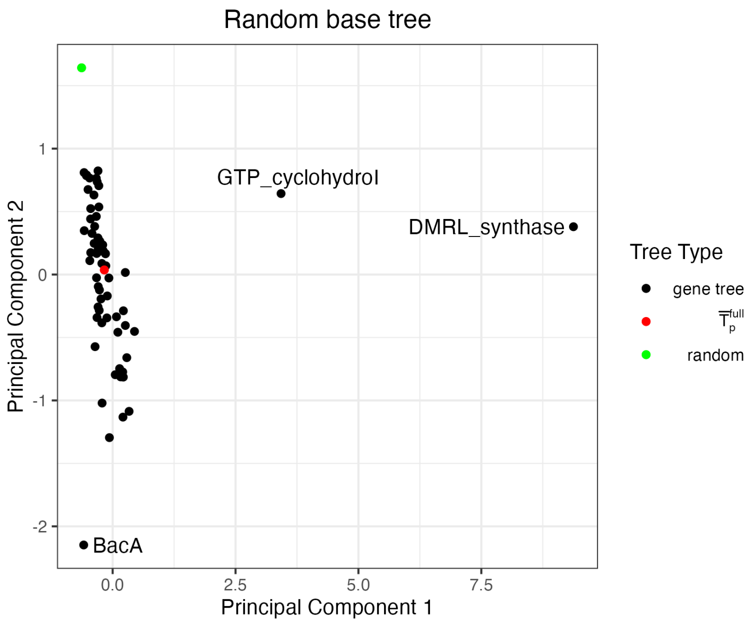


Figure 3: The proposed visualization of 63 gene trees constructed from 78 genomes from the *Prevotella* genus, depicted by a two-dimensional scatterplot. The base tree is a randomly generated tree with 78 tips.

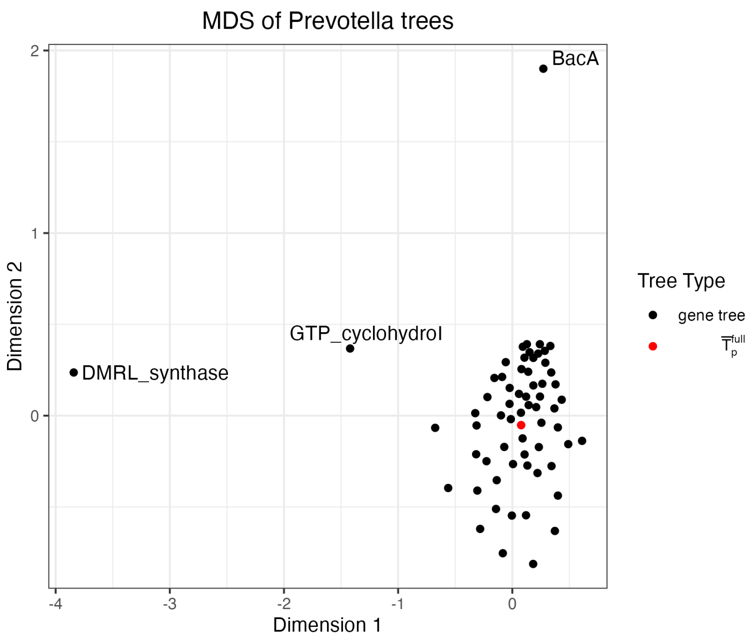


Figure 4: A visualization of 63 gene trees constructed from 78 genomes from the *Prevotella* genus using MDS of BHV distances between trees, depicted by a two-dimensional scatterplot.

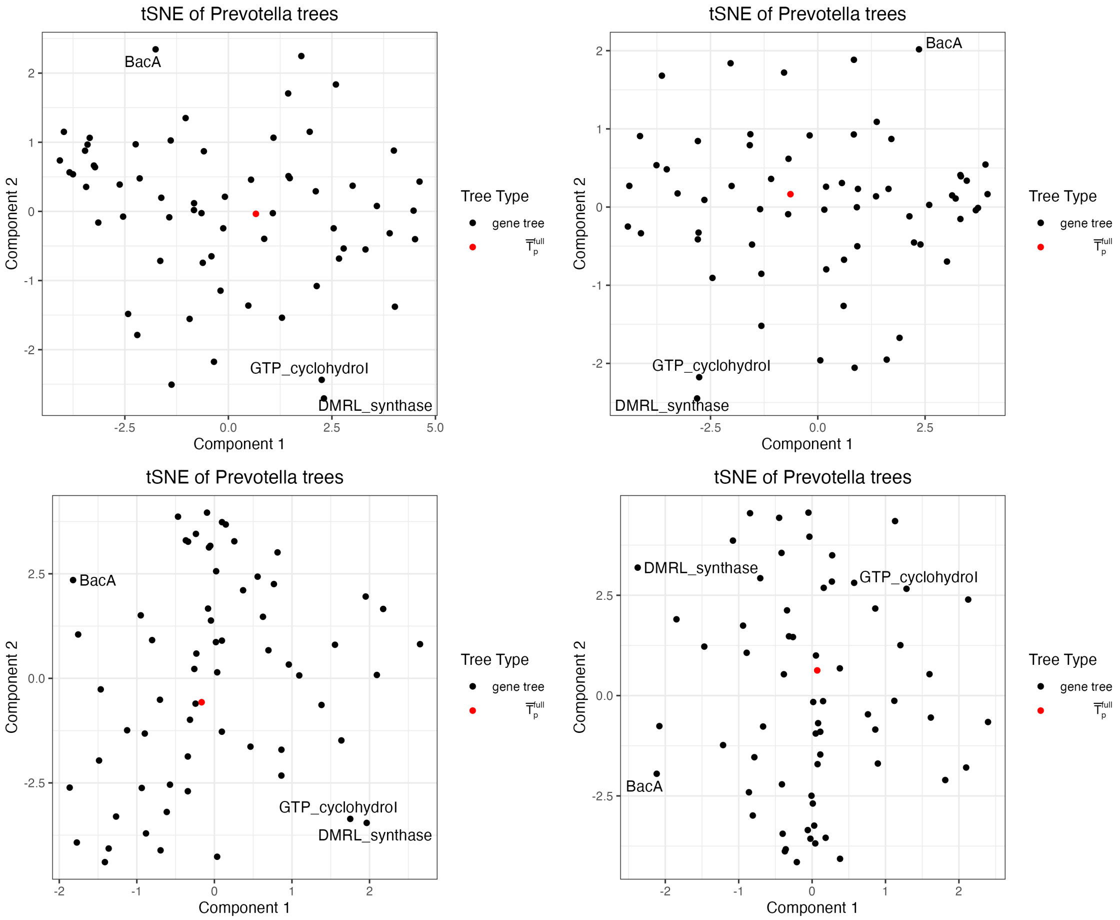


Figure 5: A visualization of 63 gene trees constructed from 78 genomes from the *Prevotella* genus, depicted by a two-dimensional scatterplot of the first two dimensions from t-SNE. Each subplot is constructed using a different initial seed. BacA, DMRL_synthase, and GTP_cyclohydroI are the outlying genes identified in the plot created with the phylogenomic tree as the base tree.

Each of the t-SNE plots show the phylogenomic tree as a central point among the set of points and BacA, DMRL_synthase, and GTP_cyclohydroI are extreme points although not separated from the other points as apparent outliers as seen in the PCA and MDS plots. The relative positions of the other points appear to be slightly different between each initial seed, although in each plot the points are spread out in a sparse pattern throughout the range of the two axes. The UMAP plots have more variation across initial seeds. The point representing the phylogenomic tree is towards the center of the points in one of the plots but is farther from the center in the other three. While the points representing BacA, DMRL_synthase, and GTP_cyclohydroI are closer to the edges of the set of points, they are not the most extreme points in the plots and would not be identified as apparent outliers. Additionally the overall pattern of the other points varies between plots. It is interesting that the plots for this analysis change so much across initial seeds and that they do not share the same outliers as the PCA and MDS plots. This suggests that different dimension reduction tools will provide different insights and may be more or less useful depending on the data being analyzed.

##### 1.4 Concordance between $\bar{T}_p^{\text{full}}$ and gene trees

Finally, we investigate the concordance between the Prevotella phylogenomic tree $\bar{T}_p^{\text{full}}$ and the gene trees. While our low-dimensional visualizations of the collection of gene trees may suggest that there is good clustering of gene trees around the phylogenomic tree, there is actually very low agreement between many of the splits in $\bar{T}_p^{\text{full}}$ and the splits present in the gene trees. We contrast the bootstrap support for $\bar{T}_p^{\text{full}}$ (computed using Hoang et al. [2018]) against the percentage of gene trees containing each split in Figures 7a and 7b. We observe the “tree of tips" phenomenon

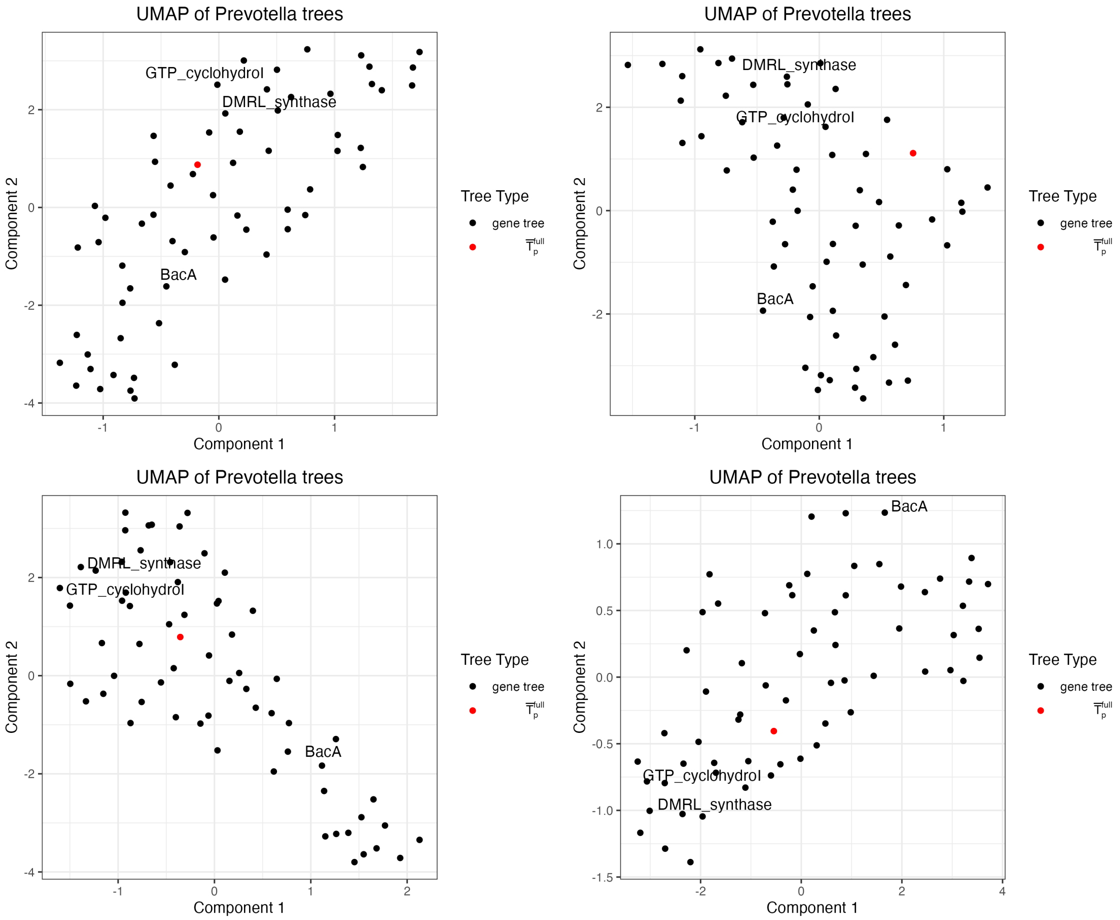


Figure 6: A visualization of 63 gene trees constructed from 78 genomes from the *Prevotella* genus, depicted by a two-dimensional scatterplot of the first two dimensions from UMAP. Each subplot is constructed using a different initial seed. BacA, DMRL_synthase, and GTP_cyclohydroI are the outlying genes identified in the plot created with the phylogenomic tree as the base tree.

described by Thiergart et al. [2014], in which splits near the tips on the tree have higher gene tree support and splits near the center of the tree have lower support. However, as noted by Avni and Snir [2020], due to high rates of HGT, a difference between individual gene trees and phylogenomic trees is to be expected. This phenomenon indicates that while the estimated gene trees agree with the phylogenomic tree for the edges that split a small number of genomes, there is much more discordance among gene trees for splits that separate larger clusters of genomes from each other. The low gene tree support for many branches on the tree contrasts starkly with the very high bootstrap support for many of these splits. As a result, we advise caution when interpreting high bootstrap support values on phylogenomic trees, as they do not necessarily imply that the individual gene phylogenies support these splits.

### 2 Streptococcus

#### 2.1 Sensitivity to base tree in log map transformation

Here we investigate the sensitivity of the proposed *Streptococcus* visualization to changes in the base tree used in the log map transformation. Figure 8 shows the proposed visualization constructed using the phylogenomic tree as the base tree (which in this analysis is also the tree with the lowest mean squared BHV distance from the other trees in the set) as well as the visualization constructed with five different base trees. Three of these trees (MnmE_helical, NFACT_N, and NAPRTase) have the smallest mean squared BHV distance from other trees in the set (Ribosomal_L9_C, Ribosomal_S30AE, and DUF3270 also have two of the smallest mean squared BHV distances from other trees but are non-binary and therefore cannot be base trees in the log map transformation). The other two trees (EcsB and DUF1934)

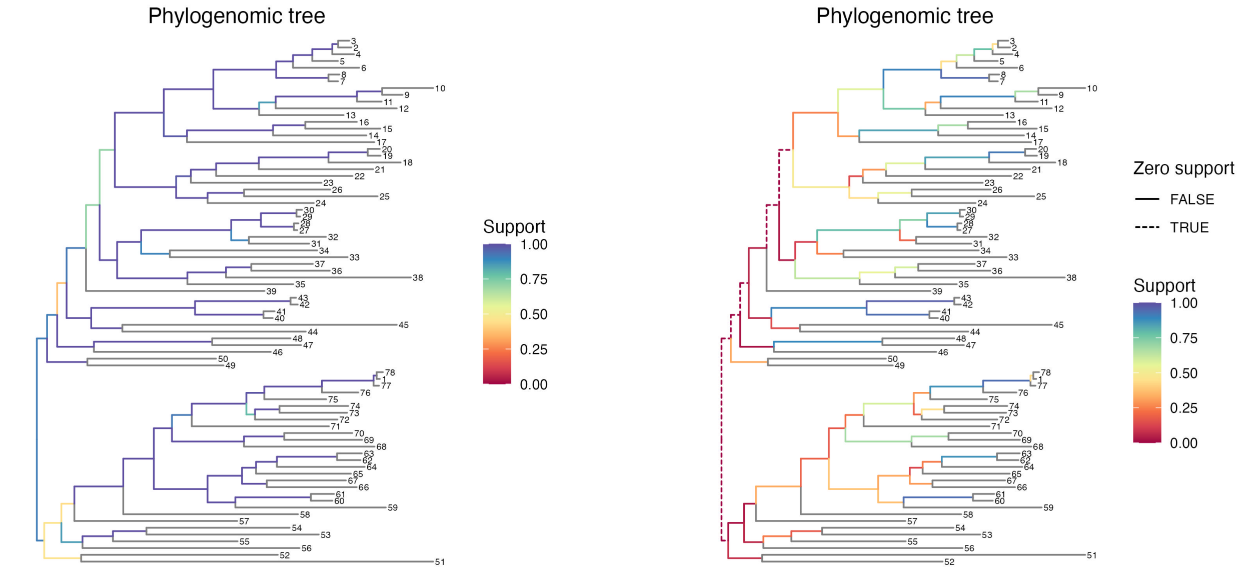


(a) Bootstrap support (b) Gene tree support

Figure 7: The Prevotella phylogenomic tree with splits colored by bootstrap support (left) and gene tree support (right). Bootstrap support is high for all splits towards the tips and for most splits farther from the tips. Gene tree support is high for splits near the tips and lower throughout the rest of the tree. We observe zero gene tree support for several splits farthest from the tips.

are apparent outliers in the visualization with the phylogenomic tree as base. While there are subtle differences between the plots, including the orientation of the first principal component, they overall show the same relationships between trees, including the dense clustering of ribosomal gene trees. This analysis appears to be robust to the choice of base tree, as trees central to the set in terms of BHV distance and trees on the edges of the visualization across all choices of base tree give similar results.

We also investigate the result of choosing a base tree with a random topology. Figure 9 shows the proposed visualization using a base tree with a randomly generated topology on 106 tips and branches drawn from the empirical distribution of non-zero branch lengths from trees in this analysis. The branch lengths must be non-zero in order for the tree to be binary and a viable base tree. As in the *Prevotella* exploration, the random base tree is an outlier (here in the first principal component) but the other patterns seen in Figure 8 can still be seen in the plot. This includes the clustering of points representing ribosomal vs non-ribosomal trees, the cone-shaped pattern of points, and the more extreme points. This provides another case in which the proposed visualization is robust even to base trees with topologies that are uninformed by the trees in the analysis.

#### 2.2 MDS visualization

Figure 10 provides a visualization of the same set of *Streptococcus* gene trees and phylogenomic tree using MDS of BHV distances. While there is a similar high density of trees in part of the plot (representing ribosomal gene trees) and the other trees are more spread out, there is a different overall shape than in the log map and PCA plots. The points are less dense than in the log map and PCA plots and collectively have a more circular pattern. While the identified outliers are at the edge of the cloud of trees, they are not easily discernible as outliers.

#### 2.3 tSNE and UMAP visualizations

As in the *Prevotella* exploration, we also create visualizations of the *Streptococcus* genomes by performing t-SNE and UMAP on the log map vectors. We use four initial seeds for these visualizations. The t-SNE visualizations can be seen in Figure 11 and the UMAP visualizations in Figure 12. Both sets of trees show the ribosomal trees clustered together in one area on the outer area of the plot, the phylogenomic tree farther from the ribosomal trees and towards the center of the cluster of trees, and the points representing genes EcsB, DUF1934, and DUF3270 on the edges of the plot. However, the overall pattern of points in the plots are different. Three of the t-SNE plots have points spread

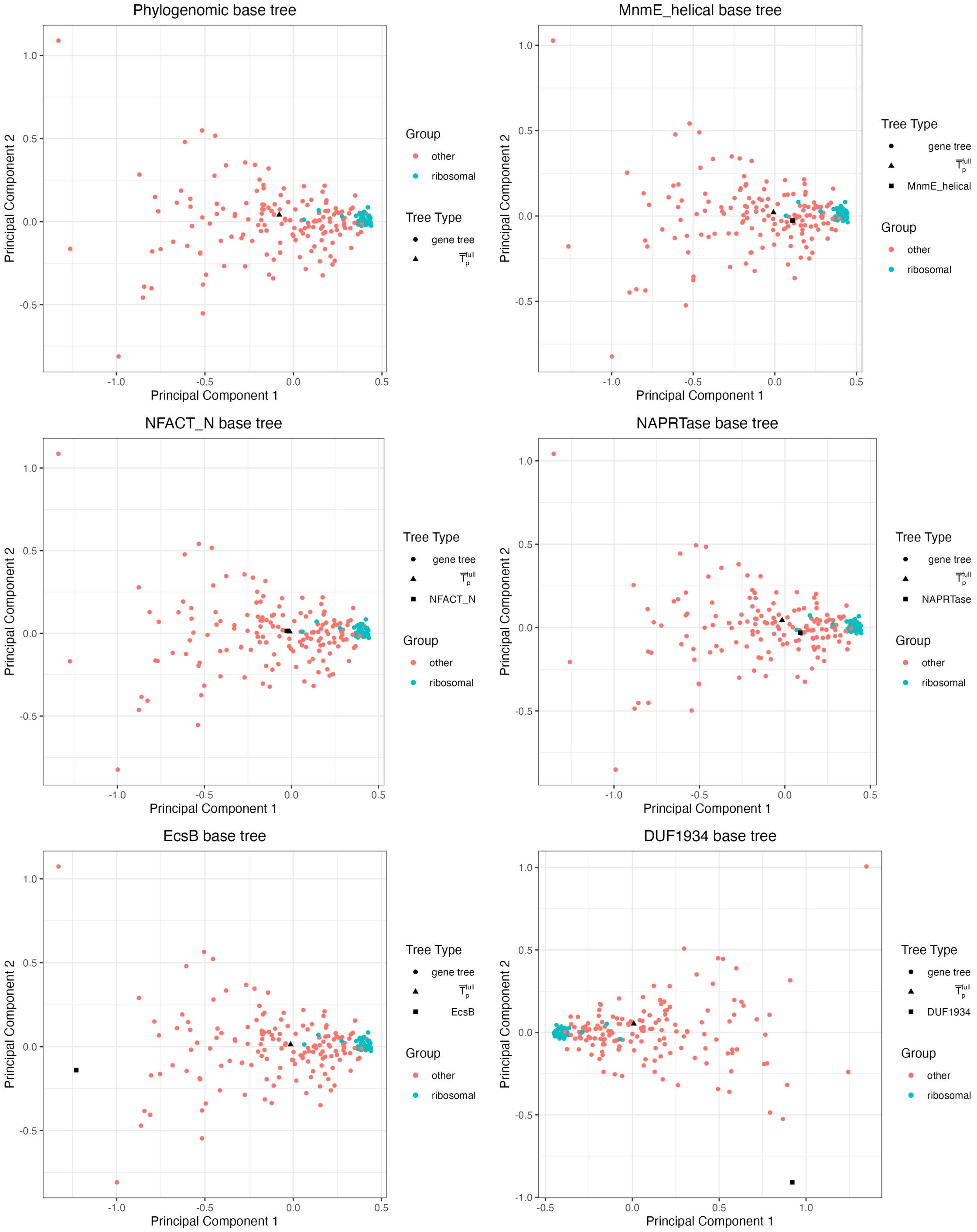


Figure 8: The proposed visualization of 196 gene trees estimated from 106 genomes in the *Streptococcus* genus shown as a two-dimensional scatterplot. The points are colored by whether or not they represent a ribosomal gene. Each subplot is constructed using a different tree as the base tree in the log map. Gene trees MnmE_helical, NFACT_N, and NAPRTase have smallest mean squared BHV distances from other trees in the set respectively out of binary trees (non-binary trees are ignored because they cannot be used as the base tree in the log map). EcsB and DUF1934 are outliers identified in the visualization with the phylogenomic tree as the base tree.

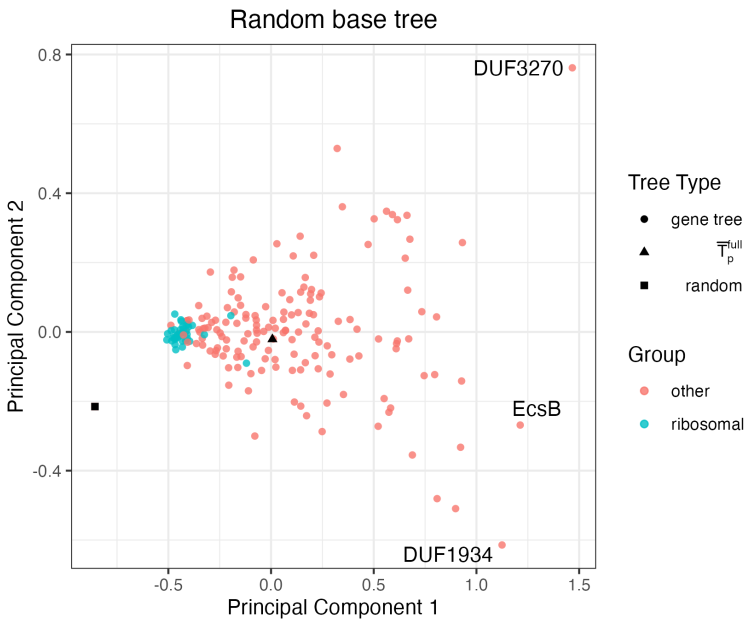


Figure 9: The proposed visualization of 196 gene trees estimated from 106 genomes in the *Streptococcus* genus shown as a two-dimensional scatterplot. The base tree is a randomly generated tree with 106 tips. The points are colored by whether or not they represent a ribosomal gene.

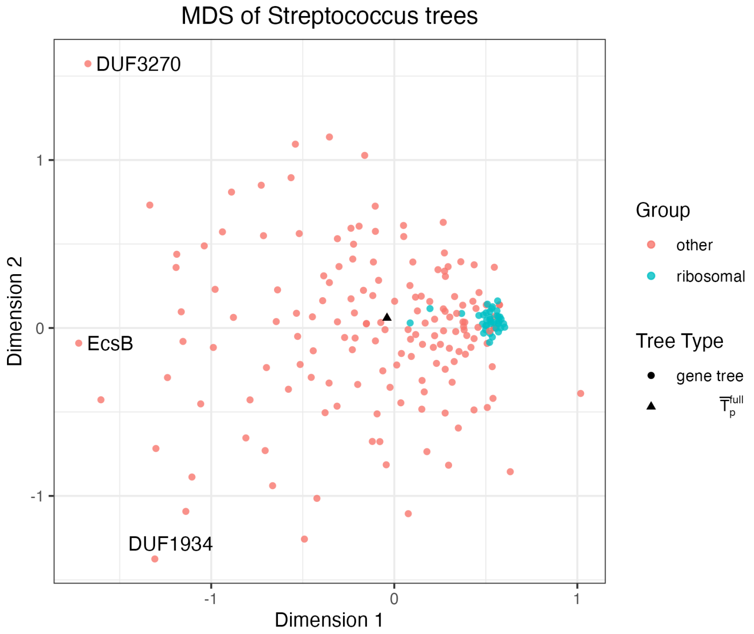


Figure 10: A visualization of 196 gene trees constructed from 106 genomes from the *Streptococcus* genus using MDS of BHV distances between trees, depicted by a two-dimensional scatterplot. The points are colored by whether or not they represent a ribosomal gene.

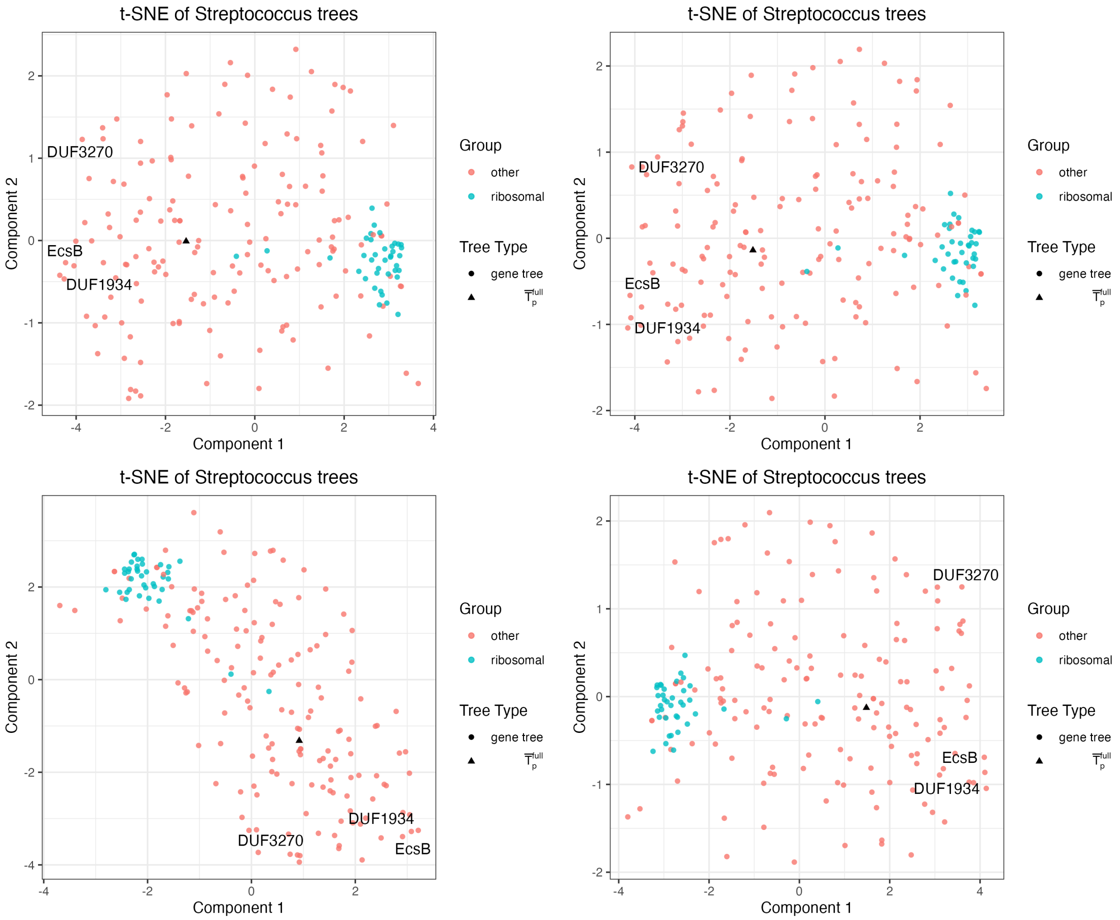


Figure 11: A visualization of 196 gene trees constructed from 106 genomes from the *Streptococcus* genus, depicted by a two-dimensional scatterplot of the first two dimensions from t-SNE. Each subplot is constructed using a different initial seed. The points are colored by whether or not they represent a ribosomal gene.

out throughout the two axes while one has more of a diagonal band of points. Conversely, three of the UMAP plots have clearly defined diagonal bands of points while one has the points more spread out throughout the two axes. This exploration provides an example in which the four different dimension reduction techniques (metric MDS of BHV distances and PCA, t-SNE, and UMAP on log map vectors) provide similar insights, although the points themselves are positioned differently.

#### 2.4 Classification using log map vectors

Once the log map transformation is used to represent a set of trees with $m$ tips as vectors in $\mathbb{R}^{2m+3}$, PCA is not the only method that can be used to explore the set of data. Consider the *Streptococcus* dataset, which includes 38 ribosomal genes and 158 non-ribosomal genes. If instead only a fraction of the genes were labeled as ribosomal or non-ribosomal, and a goal of the analysis was to predict the class of the non-labeled genes, this could be addressed with a supervised method. To illustrate this idea, we use Fisher's discriminant analysis and Random Forests to construct classifiers for the log map vectors of the gene trees, using the log map vectors from 148 trees with class labels as the training set and the log map vectors from 49 trees with hidden class labels as the test set.

We run discriminant analysis with the R function `lda` from the package MASS [Venables and Ripley, 2002]. This classifier has a prediction success rate of 63% on the test set. We also do classification with the R function `randomForest` from the package of the same name [Liaw and Wiener, 2002], which has a prediction success rate of 94% on the test set. Additionally, we are able to look at variable importance measures from our Random Forest to see which log map vectors have the largest impact on the classification algorithm. The most influential vectors are those that correspond

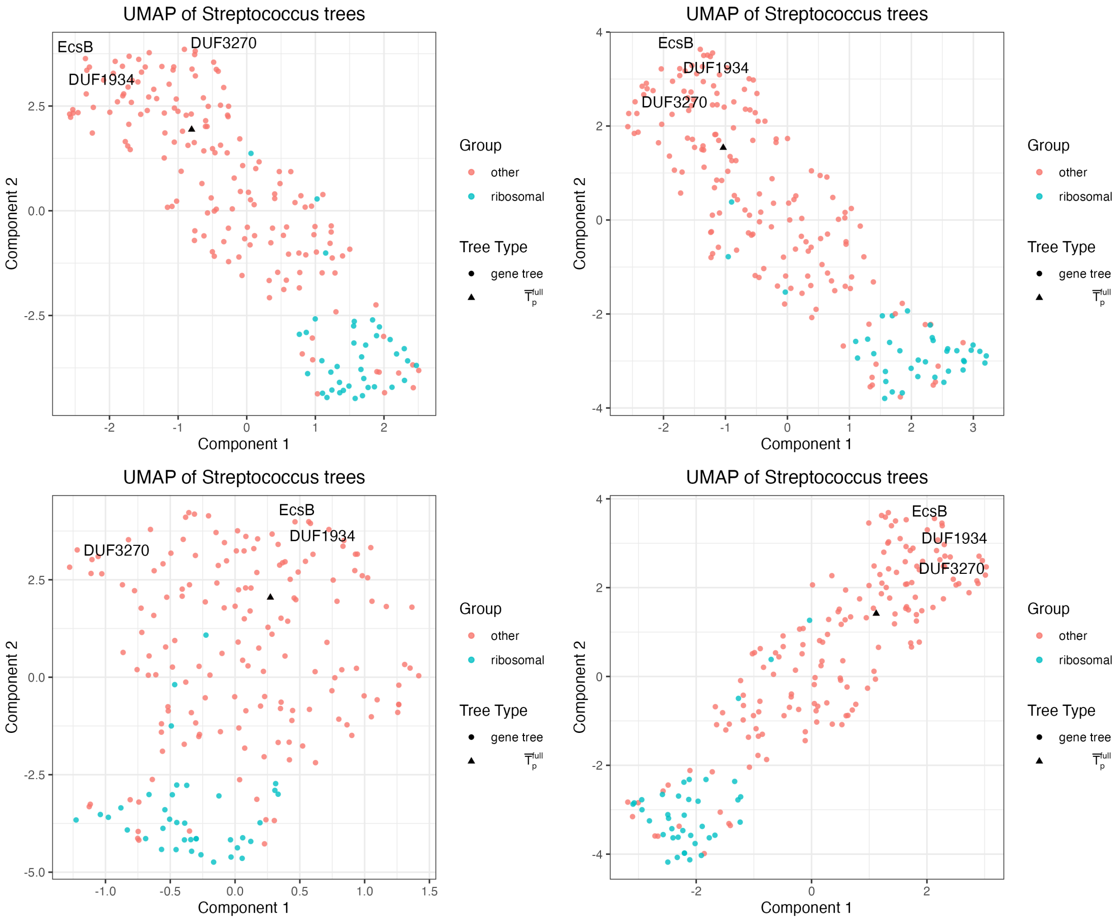


Figure 12: A visualization of 196 gene trees constructed from 106 genomes from the *Streptococcus* genus, depicted by a two-dimensional scatterplot of the first two dimensions from UMAP. Each subplot is constructed using a different initial seed. The points are colored by whether or not they represent a ribosomal gene.

with edges 100, 88, 205, and 57 in the phylogenomic tree used as base tree. These edges are shown in green, purple, red, and blue respectively in Figure 13.

#### 3 Anomaly detection in a different *Prevotella* analysis

In Section 3 we illustrated the proposed approach by investigating the robustness of the phylogenomic tree estimate to genes with apparently outlying phylogenies, and to the functional constraints on genes used to estimate the phylogenomic tree. In addition, our tool can also be used to detect anomalies in upstream processing of microbial sequence data. For example, when applying this tool to a set of 63 *Prevotella* genomes (a distinct set of genomes from those analyzed in Section 3.1), we found that the phylogenomic tree itself appeared as an outlier, and had a median gene tree support value of 0.00 (Figure 14a). We could not explain this highly unusual finding until we re-ran our entire analysis and this time estimated a dramatically different phylogenomic tree and log map visualization (Figure 14b). The second analysis showed a much more typical placement of the phylogenomic tree amongst the gene trees, and the new phylogenomic tree had a median gene tree support value of 0.40. IQTREE2's search for the maximum likelihood tree is stochastic, and in this case failed to produce a good estimate on the first attempt with the settings that we used.

We believe that the choice of base tree will have a relatively small impact on our proposed visualization in most settings, because of the similarity in topology between all trees in an analysis. This is not the case in this example because the base tree used in the first estimation run has a notably different topology from the gene trees in the analysis (this can be seen from the fact that the mean RF distance from trees in the analysis for the base tree in the first estimation run is 108, versus 66 in the second estimation run). This example demonstrates a situation in which the visualization is sensitive to

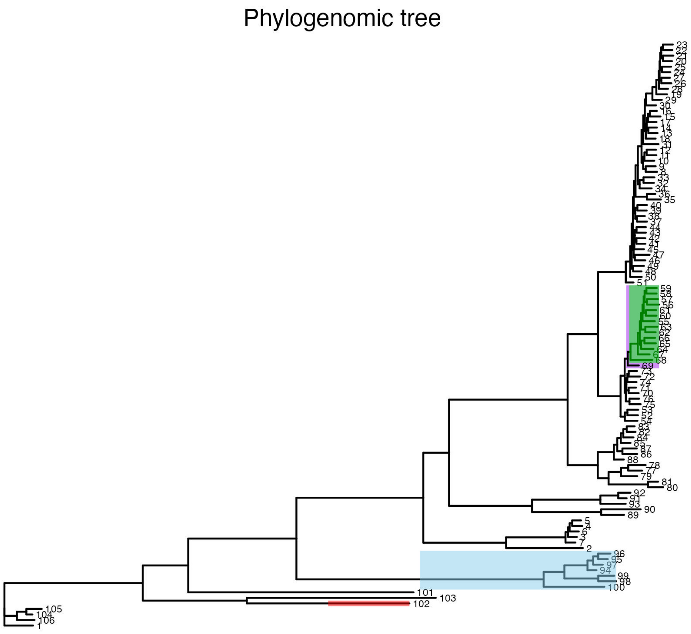


Figure 13: A phylogenomic tree for 106 genomes from the *Streptococcus* genus. The edges separating colored clades from the rest of the tree represent the edges with the highest variable importances in the Random Forest used to predict whether a gene tree is ribosomal or not.

base tree choice and there is a greater than usual difference in topology between the base tree and set of gene trees, unlike the situations shown in Appendix Sections 1.1 and 2.1.

This exploration highlights how our tool can be additionally useful for diagnosing anomalies in upstream bioinformatic and statistical analyses. We can also see the effect of choosing a base tree that has a large difference in topology from the set of gene trees. It is notable that the BacA gene is an apparent outlier in the set of trees from two distinct analyses of *Prevotella* genomes.

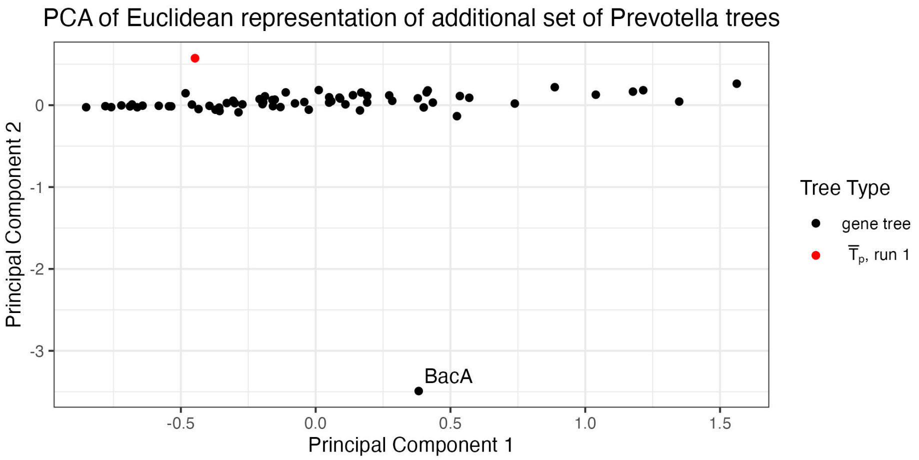


(a) First estimation run

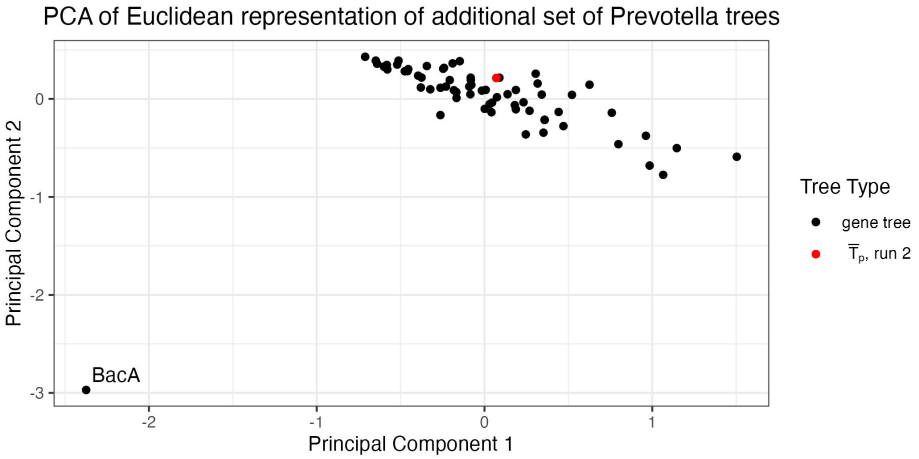


(b) Second estimation run

Figure 14: The proposed visualization of 65 gene trees constructed from 63 genomes from the *Prevotella* genus and a phylogenomic tree constructed using all genes. The two phylogenomic trees shown in red in both the top and bottom panels were estimated from two different runs of IQTREE2. The placement of the estimated phylogenomic tree on the visualization differs substantially between the two runs.
